# Statistical power of clinical trials has increased whilst effect size remained stable: an empirical analysis of 137 032 clinical trials between 1975-2017

**DOI:** 10.1101/225730

**Authors:** Herm J. Lamberink, Willem M. Otte, Michel R.T. Sinke, Daniël Lakens, Paul P. Glasziou, Joeri K. Tijdink, Christiaan H. Vinkers

**Affiliations:** Department of Child Neurology, Brain Center Rudolf Magnus, University Medical Center Utrecht and Utrecht University, P.O. Box 85090, 3508 AB, Utrecht, The Netherlands; Biomedical MR Imaging and Spectroscopy group, Center for Image Sciences, University Medical Center Utrecht and Utrecht University, Heidelberglaan 100, 3584 CX Utrecht, The Netherlands; School of Innovation Sciences, Eindhoven University of Technology, Den Dolech 1, 5600 MB Eindhoven, The Netherlands; Centre for Research in Evidence-Based Practice, Faculty of Health Sciences and Medicine, Bond University, Gold Coast, Queensland, Australia; Department of Philosophy, VU University, De Boelelaan 1105, 1081 HV Amsterdam, The Netherlands; Department of Psychiatry, Brain Center Rudolf Magnus, University Medical Center Utrecht and Utrecht University, Heidelberglaan 100, 3584 CX Utrecht, The Netherlands

**Keywords:** statistical power, clinical trial, randomized

## Abstract

**Background:** Biomedical studies with low statistical power are a major concern in the scientific community and are one of the underlying reasons for the reproducibility crisis in science. If randomized clinical trials, which are considered the backbone of evidence-based medicine, also suffer from low power, this could affect medical practice.

**Methods:** We analysed the statistical power in 137 032 clinical trials between 1975 and 2017 extracted from meta-analyses from the Cochrane database of systematic reviews. We determined study power to detect standardized effect sizes according to Cohen, and in meta-analysis with p-value below 0.05 we based power on the meta-analysed effect size. Average power, effect size and temporal patterns were examined.

**Results:** The number of trials with power ≥80% was low but increased over time: from 9% in 1975–1979 to 15% in 2010–2014. This increase was mainly due to increasing sample sizes, whilst effect sizes remained stable with a median Cohen’s h of 0.21 (IQR 0.12-0.36) and a median Cohen’s d of 0.31 (0.19-0.51). The proportion of trials with power of at least 80% to detect a standardized effect size of 0.2 (small), 0.5 (moderate) and 0.8 (large) was 7%, 48% and 81%, respectively.

**Conclusions:** This study demonstrates that sufficient power in clinical trials is still problematic, although the situation is slowly improving. Our data encourages further efforts to increase statistical power in clinical trials to guarantee rigorous and reproducible evidence-based medicine.

## Introduction

The practice of conducting scientific studies with low statistical power has been consistently criticized across academic disciplines (1–5). Statistical power is the probability that a study will detect an effect when there is a true effect to be detected. Underpowered studies have a low chance of detecting true effects and have been related to systematic biases including inflated effect sizes and low reproducibility (6, 7). Low statistical power has been demonstrated, amongst others, in the fields of neuroscience and psychology (4, 8, 9). For clinical trials in the field of medicine, the issue of sample size evaluation and statistical power is essential since clinical decision making and future research are based on these clinical trials (10, 11). Moreover, low power in clinical trials may be unethical in light of the low informational value from the outset while exposing participants to interventions with possible negative (side) effects (1). Also in medical research statistical power is low (3, 8), but a systematic overview of temporal patterns of power, sample sizes, and effect sizes across medical fields does not exist. In the current study, we provide a comprehensive overview of study power, sample size, and effect size estimates of clinical trials published since 1975 which are included in the Cochrane database of systematic reviews, and analyse emerging trends over time.

## Materials and Methods

Data were extracted and calculated from trials included in published reviews from the second Issue of the 2017 Cochrane Database of Systematic Reviews. Cochrane reviews only include meta-analyses if the methodology and outcomes of the included trials are comparable in cross study populations. Meta-analysis data is available for download in standardized XML-format for those with an institutional Cochrane Library license. We provide open-source software to convert these data and reproduce our entire processing pipeline (15).

Trials were selected if they were published after 1974 and if they were included in a meta-analysis based on at least five trials. For each individual clinical trial, publication year, outcome estimates (odds or risk ratio, risk difference or standardized mean difference) and group sizes were extracted. The power of individual trials was first calculated for detecting small, medium and large effect sizes (Cohen’s d or h of 0.2, 0.5 and 0.8, respectively); (11), based on the sample sizes in both trial arms, using a 5% α threshold. Next, analyses were performed based on the observed effect size from meta-analyses with a p-value below 0.05, irrespective of the p-value of the individual trial; if a meta-analysis has a p-value higher than 0.05, the null-hypothesis “there is no effect” cannot be discarded, and power cannot be computed for absent effects. Given that publication bias inflates meta-analytic effect size estimates (7, 13), this can be considered a conservative approach. All analyses were carried out in R using the ‘pwr’ package (16). Following minimum recommendations for the statistical power of studies (11), comparisons with a power above or equal to 80% were considered to be sufficiently powered. Study power, group sizes and effect sizes over time were summarised and visualized for all clinical trials.

## Results

Data from 137 032 clinical trials were available, from 11 852 meta-analyses in 1918 Cochrane reviews. Of these trials 8.1% had a statistical power of at least 80% (the recommended minimum (11), which we shall denote as ‘sufficient power’) to detect a small effect (Cohen’s d or h 0.2), 48% and 81% of trials had sufficient power to detect a moderate (Cohen’s d or h 0.5) or large effect (Cohen’s d or h 0.8), respectively (Figure 1). This figure shows that there was no difference between trials included in meta-analyses with a p-value below 0.05 and those above this threshold.

**Figure 1.**
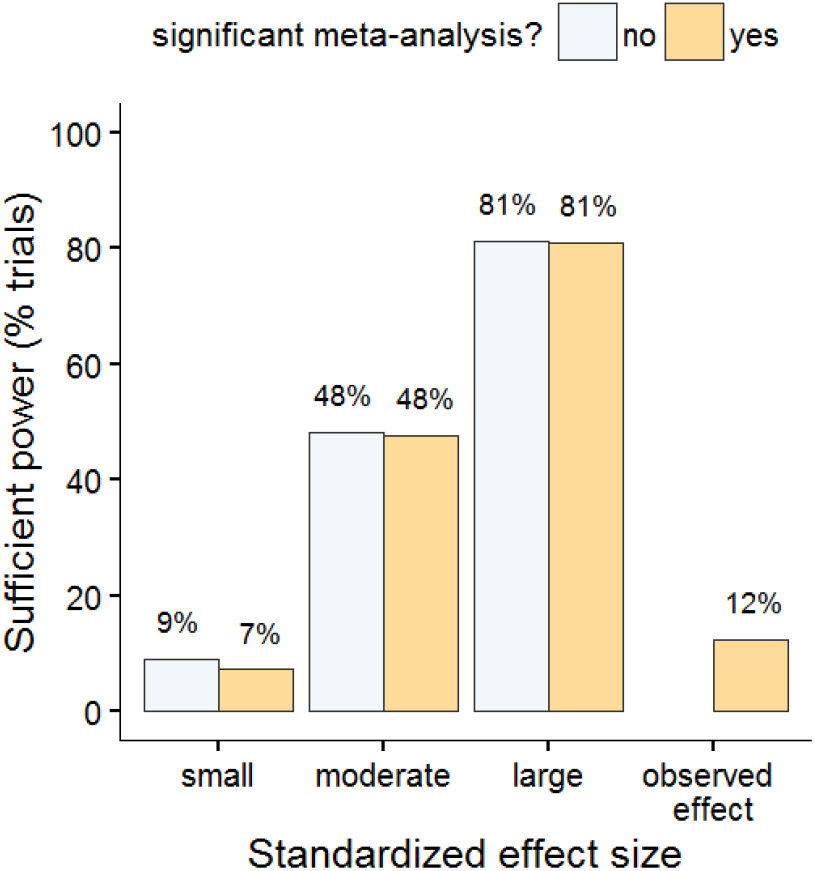
The proportion of trials that are sufficiently powered (≥80% power) for finding a hypothetical effect equal to Cohen’s d or h of 0.2 (small effect), 0.5 (moderate effect) and 0.8 (large effect) comparing trials from ‘significant’ (p-value <0.05, n = 78,401) and ‘non-significant’ (n = 58,631) meta-analyses, and the proportion of trials sufficiently powered for finding the effect as observed in the respective significant meta-analysis.

To compute study power to detect the effect observed in the meta-analysis, we examined the subset of meta-analyses with overall group differences with a p-value <0.05: 78 281 trials (57.1%) from 5903 meta-analyses (49.8%) in 1411 Cochrane reviews (73.6%). All following results are based on this sub-selection of meta-analyses. On average, 12.5% of these trials were sufficiently powered to detect the observed effect size from the respective meta-analysis (Figure 1). The median (interquartile range, IQR) power for the four categories corresponding to Figure 1 was 19% (12-37%), 78% (49-98%), 99% (87-100%) and 20% (10-48%), respectively.

Between 1975-1979 and 2010-2014 study power increased, with the median rising from 16% (IQR 10-39) to 23% (IQR 12-55) (Figure 2, left), and the proportion of sufficiently powered studies rose from 9.0% (95% confidence interval (CI) for proportions 7.6-10.6) to 14.7% (95% CI 13.9 - 15.5) (Figure 2, top right). This trend is also seen across medical disciplines (Supplementary Figure 2). When the power threshold is set at a minimum of 50% power, the proportion of trials with sufficient power is still low but also rising: from 19.3% (95% CI 17.3-21.4) in the late 1975-1979 to 27.5% (95% CI 23.5-31.9) in 2010-1014 (Supplementary Figure 1). The distribution of power showed a bimodal pattern, with many low-powered studies and a small peak of studies with power approaching 100% (Figure 2, bottom right).

**Figure 2.**
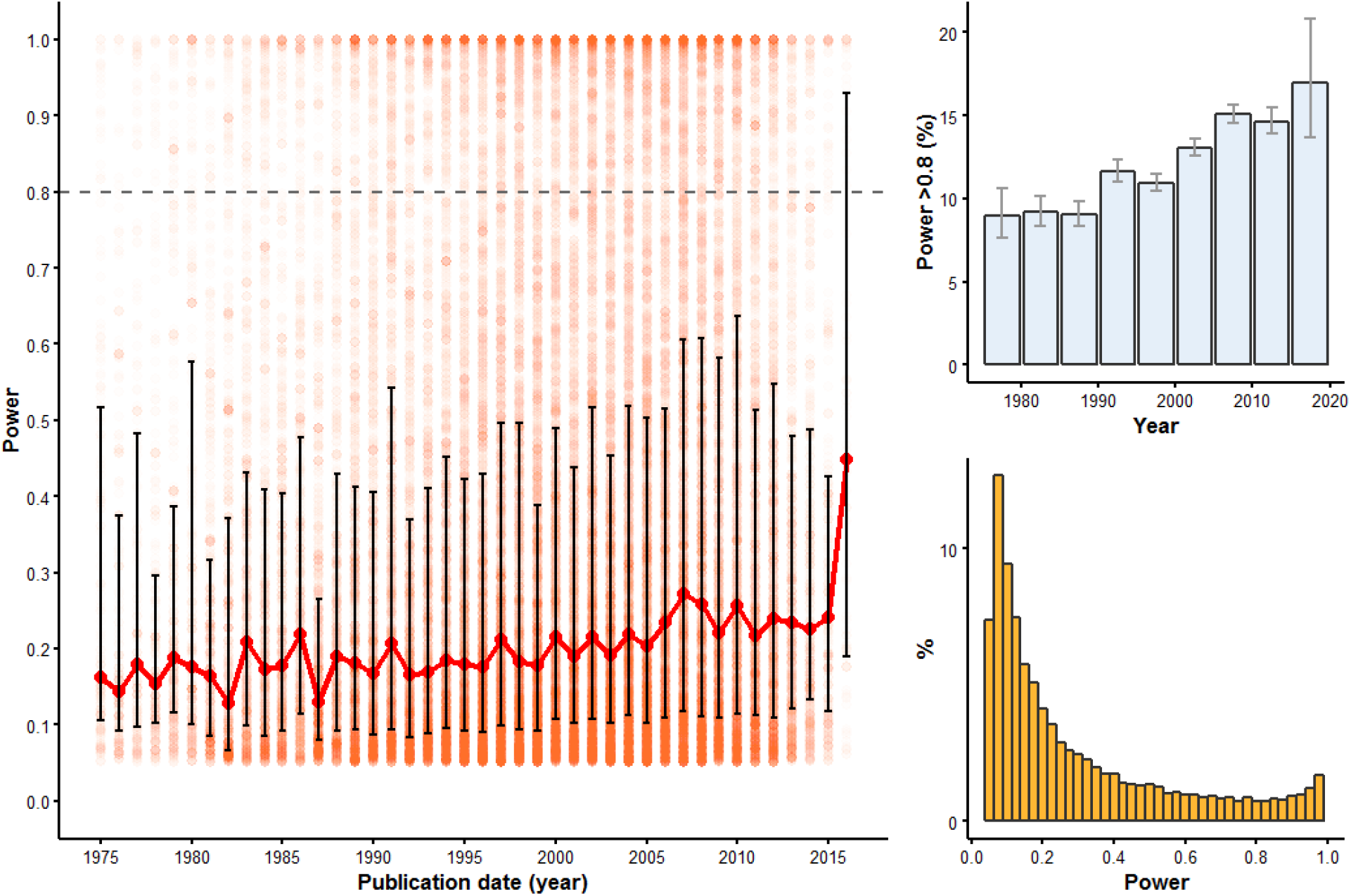
Statistical power of clinical trials between 1975 and 2017 from meta-analyses with p-value <0.05 (left). Individual comparisons are shown as semi-transparent dots. Median power is shown in red with interquartile range as error bars. The percentage of adequately powered trial comparisons (i.e. ≥80% power) is increasing over time (top right). The biphasic power distribution of the trials in general is apparent (bottom right).

The average number of participants enrolled in a trial arm increased over time (Figure 3, top left). The median group size in 1975-1979 ranged between 33 and 45; for the years 2010-2014 the median group size was between 74 and 92. The median effect sizes are summarized in Table 1; these remained stable over time (Figure 3). The standardized effect sizes were small to moderate, with a median Cohen’s h of 0.21 (IQR 0.12-0.36) and a median Cohen’s d of 0.31 (0.19-0.51) (Table 1); Supplementary Figure 3 shows the distribution plots for these two measures.

**Figure 3.**
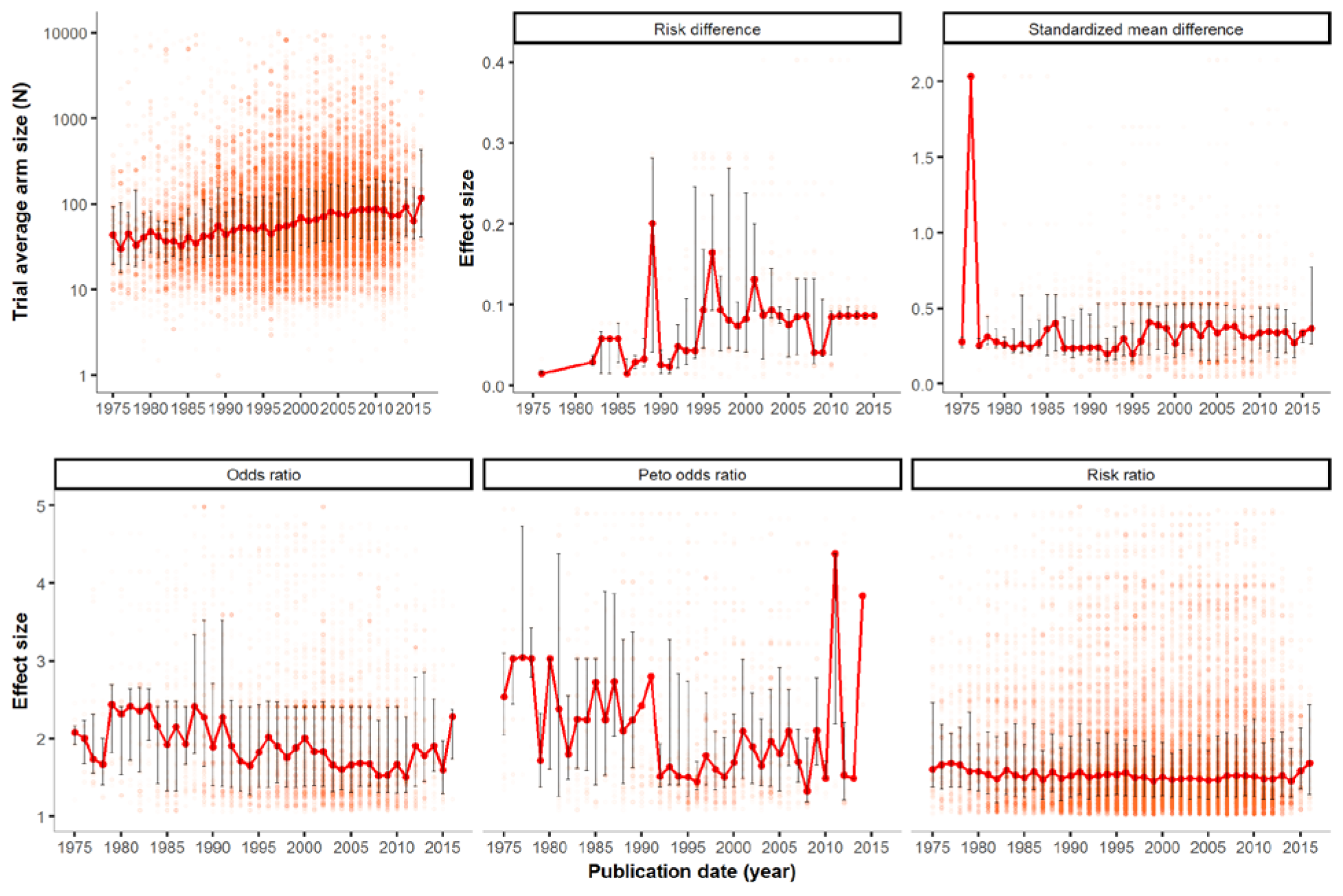
The number of participants (N) enrolled in each trial arm, between 1975 and 2017 in red semi-transparent dots (top left). Corresponding effect sizes – classified in Cochrane reviews as risk difference, standardized mean difference, (Peto) odds ratio or risk ratio – are shown in the remaining plots. Median and interquartile data are plotted annually.

**Table 1.**
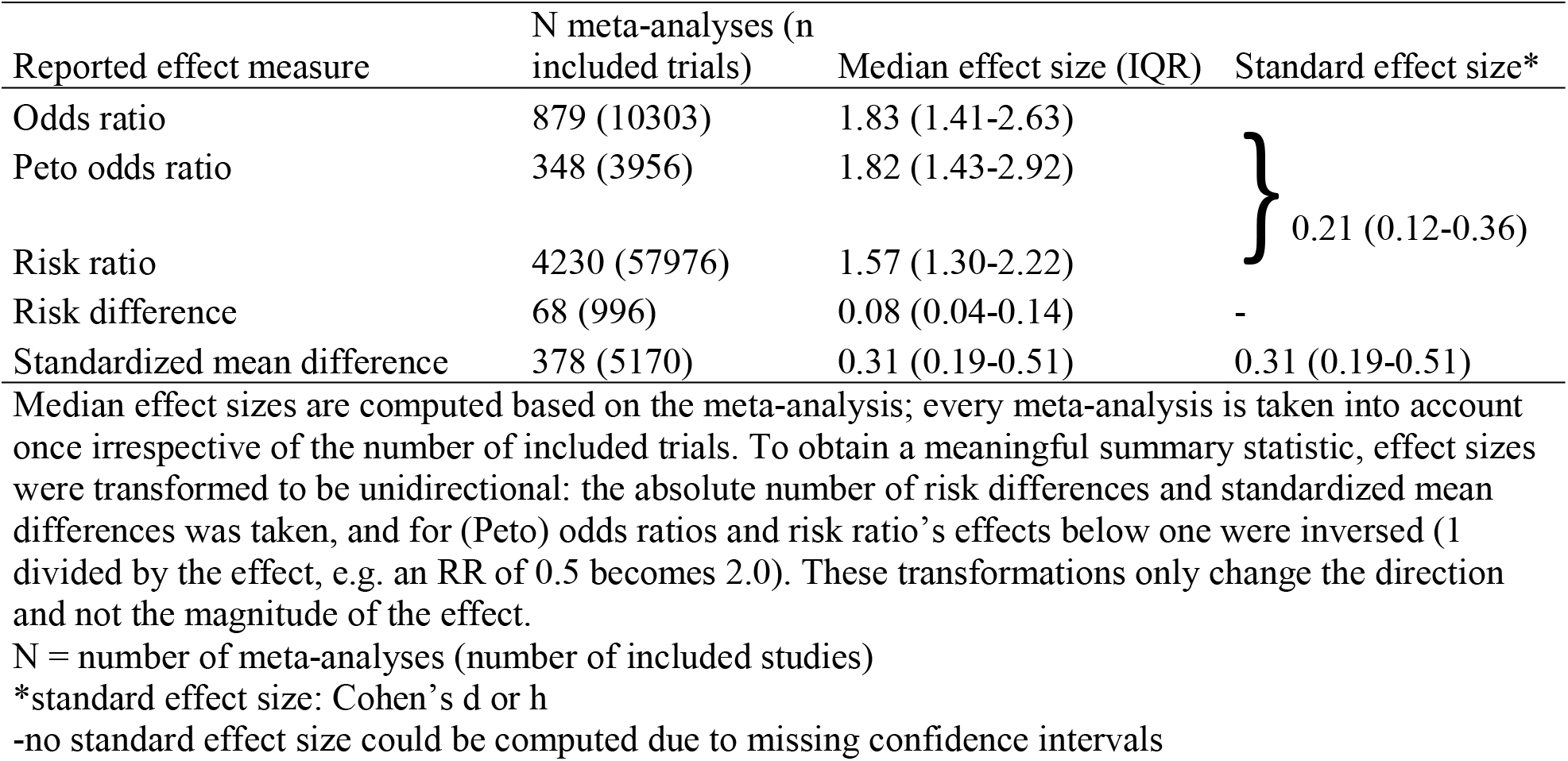
Median effect sizes for all meta-analyses with p-value <0.05

## Discussion

Our study provides an overview of the statistical power in 137 032 clinical trials across all medical fields. This analysis suggests most clinical trials are too small: the majority had insufficient power to detect small or moderate effect sizes, whereas most observed effects in the meta-analyses were actually small to moderate. Only 12% of individual trials had sufficient power to detect the observed effect from its respective meta-analysis. This study adds to the existing evidence that low statistical power is a widespread issue across clinical trials in medicine (3). Though there is considerable room for improvement, an encouraging trend is the number of trials with sufficient power has increased over four decades from 9% to 15%. On average, sample sizes have doubled between 1975 and 2017 whereas effect sizes did not increase over time.

The average effect sizes were small, with a median Cohen’s h of 0.21 and a median Cohen’s d of 0.31. These results show that large effects are rare, which should be taken into account when designing a study and determining the required minimum sample size. The effect size summary statistics provided here could also be used as standard prior in Bayesian modelling in medical research, since they are based on many thousands of trials covering the general medical field.

The study by Turner et al. (3) also used the Cochrane library (the 2008 edition) to investigate power. They showed low power of clinical trials, and a bimodal distribution of statistical power with many low-powered studies and a small peak of high-powered studies; a result also shown in neurosciences (8) and replicated by our analysis. By analysing the temporal pattern across four decades, we have been able to identify an increase of study power over time. Moreover, since effect size estimates remained stable across time, our study clearly shows the need to increase sample sizes to design well-powered studies. A study on sample sizes determined in preregistration on ClinicalTrials.gov between 2007-2010 showed that over half of the registered studies included a required sample of 100 participants or less in their protocol (12). Our findings are in line with these results, and although the average sample size has doubled since the 1970’s, we found that the median sample size in the 2010’s was still below 100.

An argument in defence of performing small (or underpowered) studies has been made based on the idea that small studies can be combined in a meta-analysis to increase power. Halpern and colleagues already explained why this argument is invalid in 2002 (1), most importantly because small studies are more likely to produce results with wide confidence intervals and large p-values, and thus are more likely to remain unpublished. An additional risk of conducting uninformative studies is that a lack of an effect due to low power might decrease the interest by other research teams to examine the same effect. A third argument against performing small studies is given in a study by Nuijten and colleagues (7), which indicates that the addition of a small, underpowered study to a meta-analysis may actually increase the bias of an effect size instead of decreasing it.

There are several limitations to consider in the interpretation of our results. First, the outcome parameter studied in the meta-analysis may be different than the primary outcome of the original study; it may have been adequately powered for a different outcome parameter. This could result in lower estimates of average power, although it seems unlikely that the average effect size of the primary outcomes is higher than the effect sizes in the Cochrane database. Second, in contrast, effect sizes from meta-analyses are considered to be an overestimation of the true effect because of publication bias (7, 13). Lastly, in determining the required power for a study a ‘one size fits all’ principle does not necessarily apply as Schulz & Grimes (14) also argue. However, although conventions are always arbitrary (11) a cut-off for sufficient power at 80% is reasonable.

With statistical power consistently increasing over time, our data offer perspective and show that we are heading in the right direction. Nevertheless, it is clear that many clinical trials remain underpowered. Although there may be exceptions justifying small clinical trials, we believe that in most cases underpowered studies are problematic. Clinical trials constitute the backbone of evidence-based medicine, and individual trials would ideally be interpretable in isolation, without waiting for a future meta-analysis. To further improve the current situation, trial pre-registrations could include a mandatory section justifying the sample size, either based on a sufficient power for a smallest effect size of interest, or the precision of the effect size estimate. Large-scale collaborations with the aim of performing a either a multi-centre study or a prospective meta-analysis might also help to increase sample sizes when individual teams lack the resources to collect larger sample sizes. Another important way to introduce long-lasting change is by improving the statistical education of current and future scientists (5).

## Key messages

- Study power in clinical trials is low: 12% of trials are sufficiently powered (≥0.8) and 23% have a power above 0.5.
- Study power has increased from 9% in 1975–1979 to 15% in 2010–2014.
- Average effect sizes are low and did not increase over time.
- When determining the required sample size of a clinical trial, moderate effects should be assumed to ensure an adequate sample size.

